# First crystal structure of double knotted protein TrmD-Tm1570 – inside from degradation perspective

**DOI:** 10.1101/2023.03.13.532328

**Authors:** Fernando Bruno da Silva, Iwona Lewandowska, Anna Kluza, Szymon Niewieczerzal, Rafał Augustyniak, Joanna I. Sulkowska

## Abstract

Herein, we present the first crystal structure of a double knotted protein TrmD-Tm1570 from *Calditerrivibrio nitroreducens*, as well the X-ray structure of each sub-domain. The protein consists of two domains TrmD and Tm1570, each embedding a single trefoil knot, which can function on their own. TrmD-Tm1570 forms a compact homodimeric complex. This protein represents one of 296 possible doubly knotted proteins from SPOUT family. Based on TrmD-Tm1570 from *Calditerrivibrio nitroreducens* we show that a double knotted protein can be fully degraded by the ClpXP degradation system, as well as its individual domains. We used numerical simulations to explain the difference in the speed of degradation. The derived kinetic parameters for the degradation process are comparable to the experimental data found for unknotted polypeptide chains.

## Introduction

Analysis of all known protein sequences has revealed the existence of proteins composed of two domains, where for each of the domains there exists a knotted crystallized homolog. This observation raised speculations that composite knots may be present in Nature. These speculations were strongly enforced by the AI predictions made by AlphaFold, which contained several structures containing composite knots, some of which were of high fidelity.^1^ Perlińska et al. put the attention towards members of the SPOUT superfamily, the most well-known group of deeply trefoil (3_1_) knotted proteins. They found four possible arrangements of domains in SPOUT family, which could lead to double knotted structures. Moreover, they showed that TrmD-Tm1570 from *Calditerrivibrio nitroreducens* (Cn) is biologically active.

Herein, we present the first crystallized double-knotted structure, based on TrmD-Tm1570 from Cn (*Cn*TrmD-Tm1570, see Fig. 1A). The protein is a member of the SPOUT superfamily,^2^ and it consists of two domains characterized as TrmD-like and Tm1570-like^3^ due to their high homology with the well known TrmD protein from *Pseudomonas aeruginosa* ^4^ and with Tm1570 from *Thermotoga maritima* ^5^ (Fig. S1 in SI Appendix).

**Figure 1:**
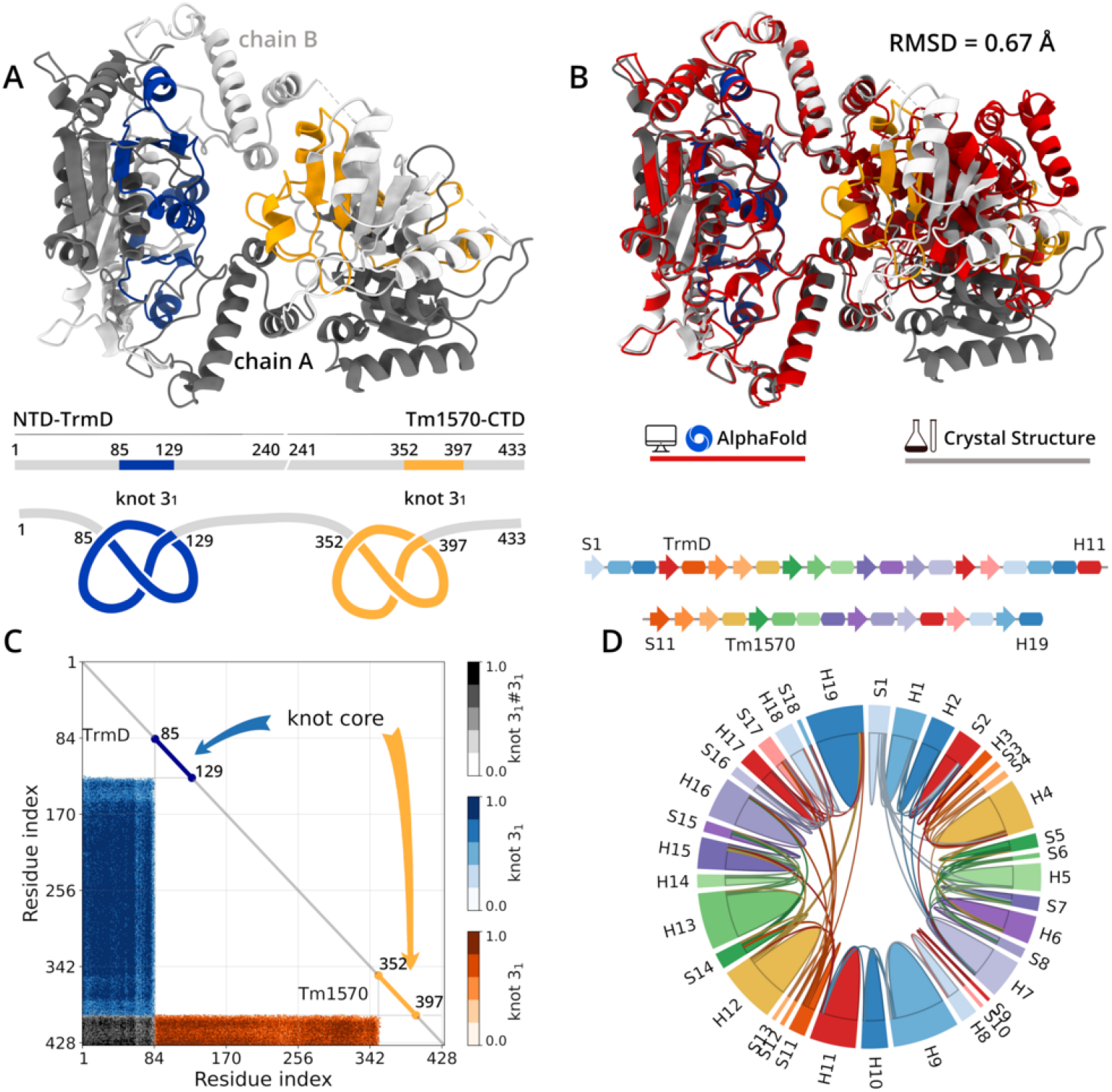
Crystal structure of TrmD-Tm1570 fusion protein from *Calditerrivibrio nitroreducens*. (A) TrmD-Tm1570 fusion homodimer complex, with chain A and B illustrated in dark and light grey color, respectively (Protein Data Bank ID code: 8B1N). The double trefoil-knot, 3_1_#3_1_, is shown in blue for NTD-TrmD and in yellow for CTD-Tm1570. The domains and knot cores positions representation of TrmD-Tm1570 fusion protein are shown in the middle of panel A. In the bottom side of panel A finds the schematic draw of double trefoil-knot. (B) Superposition of predict structure by AlphaFold at DeepMind with TrmD-Tm1570 crystal structure. The RMSD between these two structures is equal 0.67 Å. (C) The knot fingerprint with K3_1_#3_1_(8_20_)3_1_ topological notation detected in chain A from TrmD-Tm1570 fusion protein. The panel shows the knot core/location and the knot tail, residues from 1 to 84 and from 397 to 433. The color bar represents the knot probability. Only the knot with the highest probability is shown, knot cutoff > 48%. (D) Top-side panel, secondary structure representation for TrmD, from S1 to H11, and Tm1570, from S11 to H19. *β*-sheet and *α*-helix are represented by rectangles (H) and arrows (S), respectively. Bottom-side panel shows the interactions between pairs of secondary structural elements. The arc of a circle represents the secondary structure and the interactions between the corresponding elements are given by chords.

The existence of knotted proteins is important for a living cell for several aspects like folding, biological activity, and degradation, to name the key three ones. There are several hypotheses on how proteins may form twist type knots during folding process (especially for deeply knotted ones).^6,7^ In general, it was shown that proteins can self-tie *via* the twist type mechanism,^8,9^ or by so-called flipping.^10^ Analysis of the flipping mechanism lead to a more general method describing possible folding scenarios of non-twist types of knots;^11^ however, double knotted proteins were not explored yet in this context.

In contrast, much less is known about the biological role of knots. Individual study has shown that the knot provided ideal environment to conduct a chemical reaction performed by a TrmD domain. The knotted active site required different binding mode of SAM^3^ in comparison to proteins which conduct the same function but are unknotted.^12,13^

From the perspective of degradation, several *in vitro* degradation studies suggest that proteasomal mechanisms can deal with single knotted proteins. Protein degradation is a vital process which allows living cells to get rid of redundant proteins, recycle amino acids, and regulate many cellular processes.^14^ It is carried out by molecular machines called proteasomes. ClpXP is a bacterial ATP-driven molecular machine which unfolds and degrades substrate proteins. The 11-amino acid ssrA-tag (AANDENYALAA) added at the C-terminal is a recognition motif for ClpXP and makes a protein a target for ClpXP proteasome. It was shown that the trefoil knot can be easily degraded by ClpXP unless there is an obstacle along the chain (e.g. stable domain) preventing the knot from tightening. ^15,16^ Also, based on the studies on human UCHs containing gordian (5_2_) knot, it was shown that even more complex topology does not pose a problem for the degradation process. ^16–18^ The observed slow kinetics of UCHs was shown to be a consequence of high local stability, especially near the tag,^16^ which was also observed in the case of a protein with trivial topology.

Altogether, even though these processes are not fully understood for single knotted proteins, now one has to ask the question how these processes may proceed in the context of proteins with composite knots.

Herein, first we present in detail crystal structure of doubly knotted protein with Mg^2+^, and SAM – *Cn*TrmD-Tm1570. Then we focus on the degradation process and show experimental results obtained for individual domains from *Cn*TrmD-Tm1570 protein as well as the whole protein and relate them to the behavior of a well known GFP protein. Furthermore, by means of Molecular Dynamics simulations, we explain possible pathways which can lead to full degradation of the double knotted protein and the difference in the kinetics between single knotted structures. Finally, we hypothesize how double knotted protein from Spout family could fold.

## Results

### Structure of double knotted protein – *Cn*TrmD-Tm157 homodimer complex

TrmD-Tm1570 protein is crystallized as a compact homodimeric complex (Fig. 1A). The *Cn*TrmD-Tm1570 protein has 433 residues in its monomeric form, and in such a structure, N-terminal domain TrmD (NTD-TrmD) includes residues 1-240 and the C-terminal domain Tm1570 (CTD-Tm1570) comprises residues 241-433. Moreover, the determined structure adopts a globular *α*/*β* structure consisting of seven *β*-sheets and nine *α*-helices in the NTD-TrmD and the CTD-Tm1570 is composed by five *β*-sheets and eight *α*-helices. In the both proteins, NTD-TrmD and CTD-Tm1570, five-stranded *β*-sheets sandwiched by *α*-helices at both sides. The structure of *Cn*TrmD-Tm1570 has been deposited to the Protein Data Bank (https://www.rcsb.org/structure/8BN1) under accession code 8BN1. In Fig. 1B, we analyzed the superposition of crystal and AlphaFold structures. The superposition displayed a well fit to the NTD-TrmD, but to the CTD-Tm1570, slight differences are observed. However, the root-mean-square deviation (RMSD) for the whole structure has a small value, RMSD equal to 0.67 Å.

The *Cn*TrmD-Tm1570 in its monomeric form is a protein with a double-knot and to identify the knot type and core - i.e., the shortest portion of the protein backbone to form the knot type, the knotting fingerprint was calculated being the only technique to detect the knot type (Fig. 1C).^19^ The *Cn*TrmD-Tm1570 has a classification K3_1_#3_1_, which the first knot is located in NTD-TrmD between amino acids 85 and 129. The core of second knot, located in CTD-Tm1570, is within range of residues 352-397. Moreover, the individual chains adopt a so-called open conformation in which the domains do not interact with each other (Fig. 1D). Instead, in the homodimer structure, domains from one monomer interact with their counterparts from the other monomer. In the complex they adopt relative antiparallel (TrmDs) and perpendicular (Tm1570s) configurations, accordingly to orientations of specific *β*-sheets located in their trefoil knots (Fig. S2 in SI Appendix).

The crystal structure of *Cn*TrmD-Tm1570 includes Mg^2+^ ion and S-adenosylmethionine (SAM) molecule in the active region of NTD-TrmD and a SAM in the active region of CTD-Tm1570 (Fig. 2). It is well known that Mg^2+^ is necessary for TrmD activity, here we show how exactly the ion binds.^12^ The Mg^2+^ binds with a residue E115 from chain A and residues E167, E168, and E176 from chain B; SAM interacts with the dimeric interface binding with residues G112, I132, G133, F135, I137 and S169 from chain A and E168 from chain B (Fig. 2A), this stays in agreement with quantum analysis which showns that E115 is necessary for the methyl transfer. ^3^ In the CTD-Tm1570, SAM interacts with residues T356, A357, G382, F387, I402, I409, L411, and Y420 from chain A (Fig. 2B).

**Figure 2:**
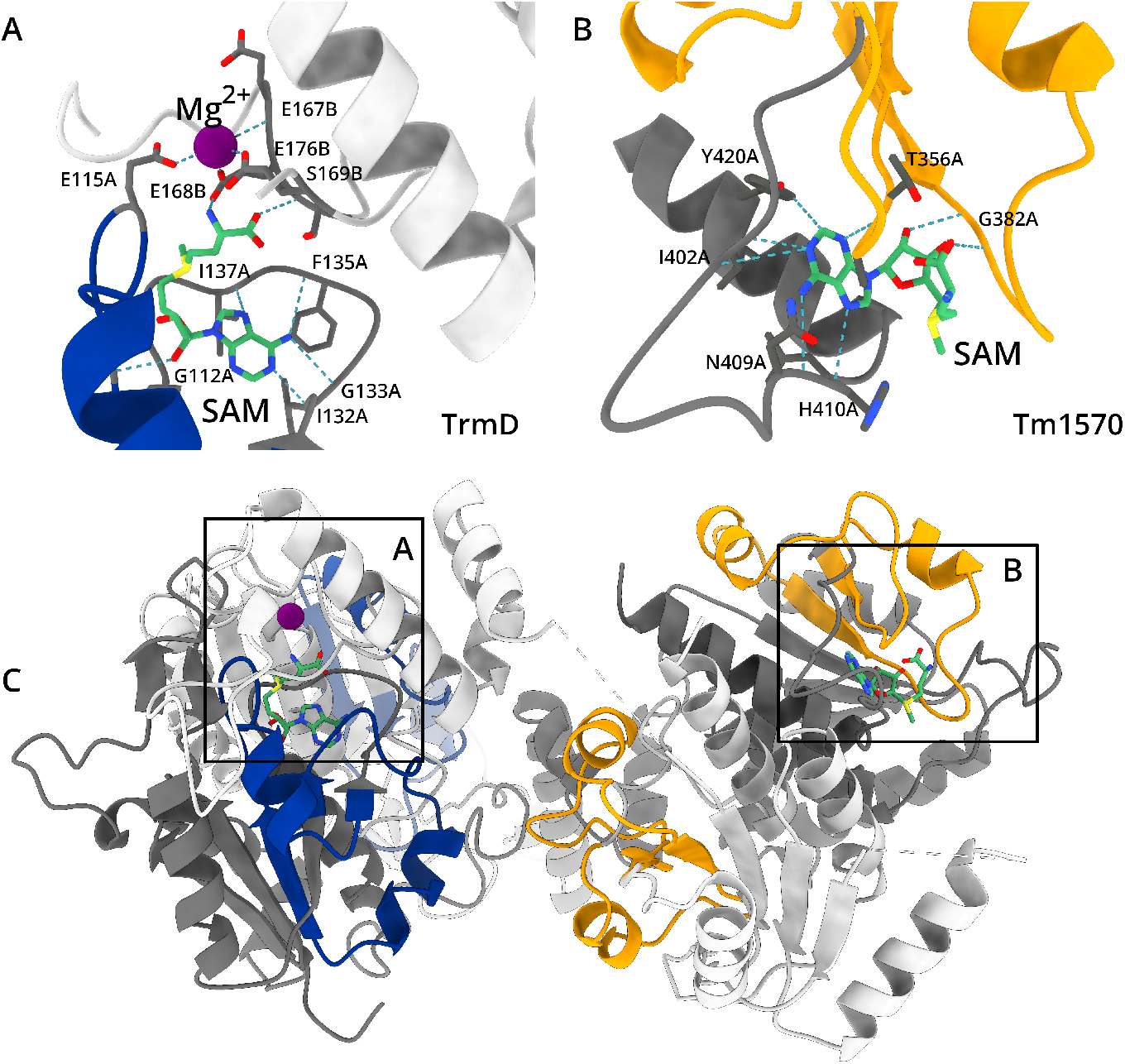
S-adenosylmethionin (SAM) and Mg^2+^ binding sites present in TrmD-Tm15780 fusion protein. (A) Dimeric interface binding between SAM (green color) and Mg^2+^ (purple color) with chain A (dark grey color) and chain B (light grey color) from NTD-TrmD. knot 3_1_ is shown in blue from chain A of NTD-TrmD. (B) Interface binding between SAM and chain A from CTD-Tm1570 (dark grey color) with its knot 3_1_ in yellow color. The amino acids in the binding sites are shown in stick representation. (C) General overview of CnTrmD-Tm1570 fusion protein (PDB ID: 8B1N) highlighting the binding sites of SAM and Mg^2+^ described in A and B.

Additionally, we were able to crystallized single TrmD from *Calditerrivibrio nitroreducens* with SAM in the binding site (PDB ID: 8BYH). Based on the superposition of *Cn*TrmD against the *Cn*TrmD-Tm1570 we found that a RMSD is just 0.68 Å across all 240 residues pairs, with (see Fig. S4 in SI Appendix). This shows that single TrmD is able to form stable structure, bind SAM and have almost the same conformation as TrmD from fusion.

In order to understand potential difference between binding SAM in fusion, and in the single domains we conducted analyses of binding SAM to TrmD and Tm1570 over different representative organisms, following our previous comprehensive analysis.^13^ We selected four TrmD+SAM from other organisms, *Haemophilus influenzae, Haemophilus influenzae Rd KW20, Pseudomonas aeruginosa UCBPP-PA14*, and *Mycobacteroides abscessus* (Fig. S3A, C, E, F, and G in SI Appendix). As predicted, the conservative interactions are those formed between SAM and the adenine-binding loop. Three H-bonds are present in all structures, to TrmD+SAM from CnTrmD+Tm1570, two interactions between NH2 (SAM) and the loop, F135A and G133A, and one interaction between ILE132A and the N atom in adenosine (SAM) (Fig. 2A). We concluded that SAM binds in the same manner to TrmD in fusion and single domain.

In the case of Tm1570 different interactions were observed between Tm1570 with SAM from the crystal structure and from the organism *Thermotoga maritimathese*. However, the interactions between SAM and its loop are still maintained, as predicted in TrmD (Fig. 2B and Fig. S3B and D in SI Appendix).

### Degradation of deeply knotted individual domains of the fusion protein

Next, we investigate degradation process *via* ClpXP proteasome of the double trefoil-knotted SPOUT protein and its both deeply knotted single-knotted domains (Tm1570-like and TrmD-like), which were also produced as individual proteins. As an additional control, we measured degradation kinetics for GFP, a protein well-studied in this context.^20,21^ proteolysis was measured by SDS/PAGE, where the progress of each reaction was determined by measuring bands intensity over time.

In order to construct proteins substrates for ClpXP, we appended ssrA-tag at the C-terminals. Both domains are degraded by ClpXP. Analysis of Michaelis-Menten steady state degradation provided following kinetic paramaters: *K*_*M*_ = 1.62 *±* 0.56 *µ*M and 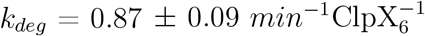 for Tm1570-ssrA and *K*_*M*_ = 4.12 *±* 1.01 *µ*M and 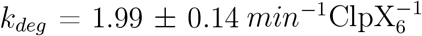 for TrmD-ssrA (Fig. 3A). For both substrate proteins we observe additional band with an increasing intensity with the progress of the reaction. This implicates an appearance of the partially degraded substrate. The estimated difference in mass of the substrate band and the intermediate band in both cases amounts to several kDa. This suggests a cut off of a few amino acids, including ssrA-tag necessary for recognition by the ClpXP complex. Such a protein cannot be absorbed and degraded by the proteasome and it manifests itself as an intermediate product.

**Figure 3:**
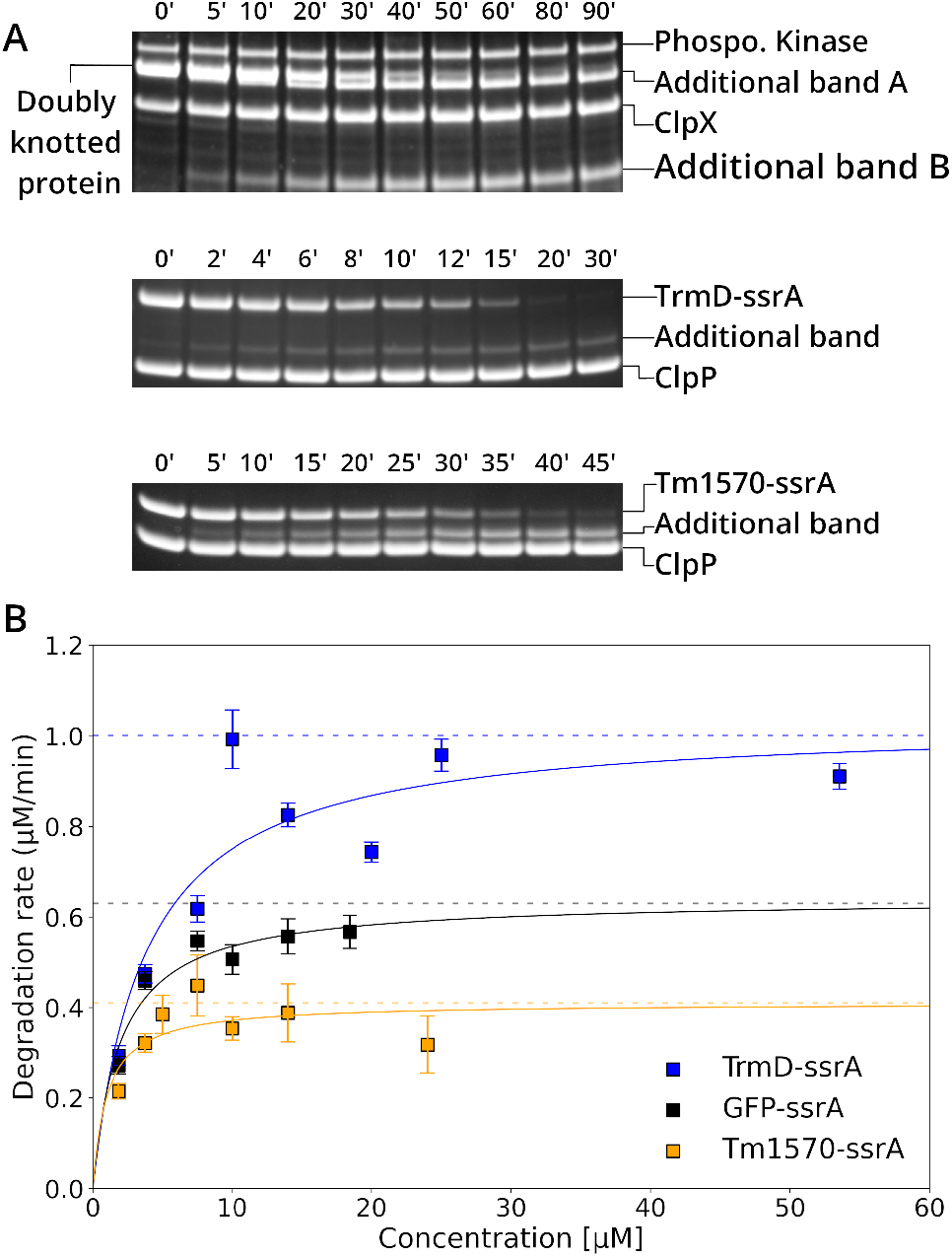
(A) upper side - The diagram of structure of doubly knotted protein; bottom side - SDS-PAGE gel for degradation of doubly knotted protein in concentrations 7.5 *µ*M. Degradation of TrmD and Tm1750, both in the same concentrations (14 *µ*M). (B) Michaelis-Menten plot of degradation for: TrmD-ssrA (green), GFP-ssrA (black), Tm1570-ssrA (red).

The degradation kinetic parameters for GFP-ssrA are equal to *K*_*M*_ = 2.22 *±* 0.61 *µ*M and 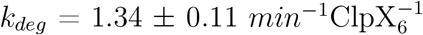. There was no addi tional band observed in the case of GFP-ssrA. TrmD-like domain is degraded faster than most of studied knotted proteins, and comparable with another deeply knotted SPOUT protein - YbeA-ssrA^16^ (Table 1 in SI Appendix). The degradation is also faster than for GFP-ssrA and degradation kinetics of Tm1570-ssrA is slower than both TrmD-ssrA domain and GFP-ssrA (Fig. 3B). However, it is comparable with kinetics observed for some stable unknotted proteins.^22,23^ These results are another confirmation that knotted proteins can be degraded using principal machinery present in a living cell.

### Degradation of double knotted protein

Following degradation of individual domains, we performed the degradation process of the whole doubly knotted protein. The densitometric analysis of the degradation reaction shows that ClpXP can degrade the substrate. The double knotted protein was degraded more slowly than the individual domains. In the case of the whole protein, the complete disappearance of the band on the gel occurred after above 60 min. For Tm1570-ssrA and TrmD-ssrA the complete decay of the band occurred on average after 15 min and 12 min, respectively. However, we were unable to determine kinetic parameters for the degradation of the substrate protein. During a gradual decrease in the intensity of a band corresponding to the protein substrate there appeared two new bands which indicate formation of partial products of the degradation process (Fig. 3A). Based on the analysis of the differences in mass on the gel, it occurs that the band A and protein substrate differ by a few kDa. The band A corresponds to the protein part that was released after being cut off by ClpXP of several dozen residues from C-terminus of the doubly knotted protein. The protein fragment that was degraded corresponds to the fragment that was degraded during Tm1570 proteolysis by ClpXP. The additional band B corresponds to a fragment of the substrate lacking the Tm1570 domain and the C-terminal TrmD domain. The band B on the gel represents the TrmD N-terminal domain that has not been degraded (similar fragment to that left after TrmD-ssrA degradation).

### Knot Degradation by Theoretical Model

To explore the degradation process of the deeply knotted protein in more detail, we applied Molecular Dynamics simulations. In this study, the coarse-grained molecular machine for protein degradation was built based on the ClpX architecture (see Fig. 4A and B). A ssrA-tag was covalently attached at the C-terminal of each knotted protein (*Cn*TrmD-Tm1570, NTD-TrmD, CTD-Tm1570, and GFP) and acted as the leading peptide to initiate the translocation process. The complex, protein and ssrA-tag, was set in the pore entrance, according to the schematic representation in Fig. 4C, to mimic the real system, in which protease recognizes a substrate.

**Figure 4:**
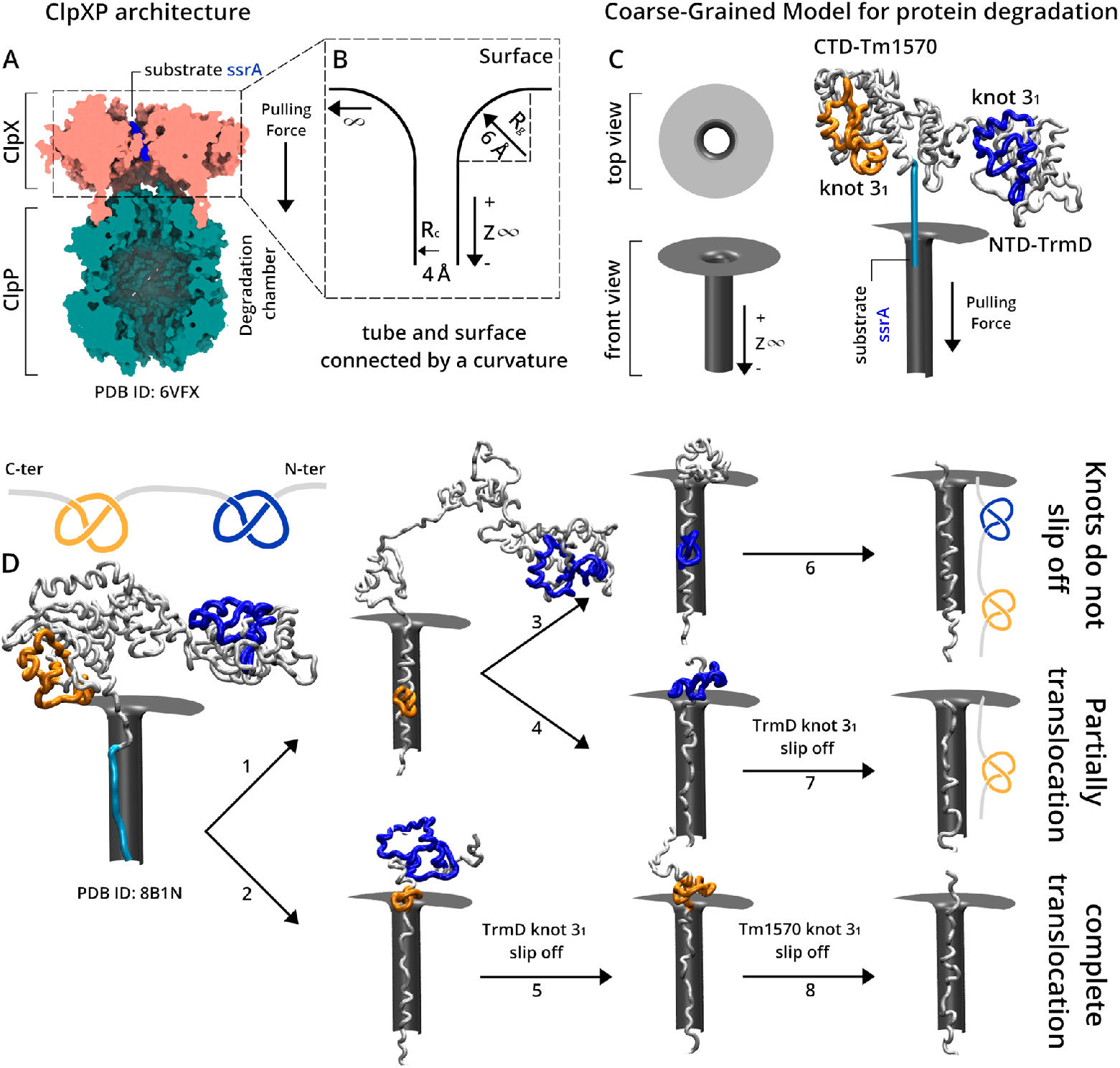
The ClpXP architecture and the coarse-grained model representation for protein degradation. (A) Cutaway view shows the axial position of the substrate ssrA (blue color), ClpX unfoldase (coral color), and ClpP protease (teal color) - degradation chamber. ClpXP structure corresponds to the PDB ID: 6VFX. (B) Schematic view of the modelled tube and surface connected by a curvature. *R*_*c*_ and *R*_*g*_ correspond to the tube radius and the curvature radius between tube and surface, respectively (See Tube Model for Protein Degradation in Materials and Methods). (C) Left-side panel, visualization of front and top view of tube and surface model with the parameters presented in panel B. Right-side panel, CnTrmD-Tm1570 fusion protein connected with ssrA by its C-terminal from CTD-Tm1570. The model was setup with ssrA inside of the tube to better reproduce the real system, panel A. (D) Snapshots from the simulation at force 22 *ϵ*/Å to CnTrmD-Tm1570 fusion protein. At low force, three degradation process was observed: Path 1 (1-3-6), complete degradation, knots do not slip off; Path 2 (1-4-7), pore obstruction by TrmD knot 3_1_, unthreding process; Path 3 (2-5-6), unthreding process observed in both knots, complete translocation from native position to N-terminal.

First, we studied the degradation process of *Cn*TrmD-Tm1570 fusion protein, which contains a double trefoil knot. The fusion protein was threaded through a narrow pore at constant force (see Fig. 4D) and the magnitude of the force during the pulling process was equal 22 *ϵ/*Å. The degradation mechanism of fusion protein has shown three distinguished degradation pathways in this specific magnitude of force. In the first case, path 1 (1-3-6), the translocation occurred to both knot cores pulled in through the pore. However, in path 2 (1-4-7), we observed in such a case that knot located in the Tm1570 region was pulled in through the pore, but the knot located in the TrmD region slipped off the chain. Lastly, in path 3 (2-5-8), a complete translocation occurred with both knots slipping off the chain. Overall, how deep the knot can be is a crucial factor affecting the translocation process and the probability of spontaneously slipping off the chain increase from shallow to depth knots. In case of individual domains, CTD-Tm1570, NTD-TrmD, and GFP, we calculated the average degradation times (Fig. 5). Degradation times were measured in a range of forces for which we could observe the knot degradation without pore obstruction. As it is shown in Fig. 5A, for forces equal to 30 *ϵ/*Å, 32 *ϵ/*Å, and 34 *ϵ/*Å, we observe similar degradation times for NTD-TrmD, GFP, and CTD-Tm1570. In such a case, forces were strong enough to break all local interactions during the pulling process and to tighten the knot in NTD-TrmD and CTD-Tm1570, and finally to complete the translocation without pore obstruction. During the translocation process, the knot size decreased and a minimal displacement was observed related to its native position.

**Figure 5:**
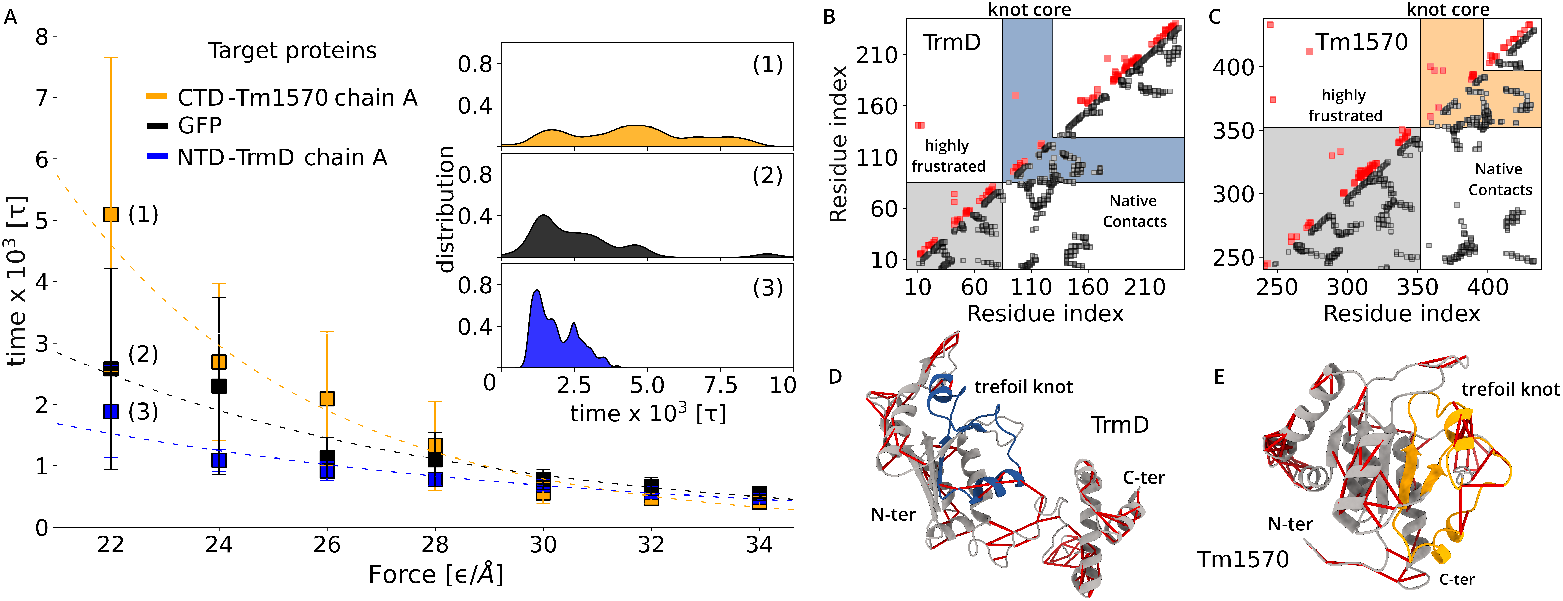
Protein Degradation of the NTD-TrmD, GFP, and CTD-Tm1570. (A) Protein degradation time [*τ*] for CTD-Tm1570 chain A, GFP, and TrmD-NTD chain in orange, black and blue colors, respectively. The degradation time was measured in a range of force [*ϵ/*Å] from 22 to 34. Right-side panel corresponds to the time distribution for the target proteins in 22 *ϵ/*Å. (B) and (C), the upper and the lower triangles are shown the highly frustrated contacts, red color, and native contacts, black color, respectively, for TrmD and Tm1570. The highlighted blue and orange regions correspond to the knot core of TrmD and Tm1570, respectively. The highlighted gray region corresponds to the distribution of the highly frustrated and native contacts which knot must go through. (D) and (E) correspond to the native structure of TrmD and Tm1570, respectively, with their highly frustrated contact distribution in red.

Differences in the degradation times among the proteins were observed for weaker force within the range from 28 *ϵ/*Å to 22 *ϵ/*Å. The difference between the degradation of these proteins become more spotlighted in this scenario. According to the experimental procedure and our theoretical method, the degradation starts from the C-terminal end. During the pulling process, specifically at force equal 22 *ϵ/*Å, in 98% of simulations for NTD-TrmD the knot slides off the chain, and before the translocation process is completed. On the other hand, for CTD-Tm1570, in only 62% of the simulations the knot slides off the chain. In the remaining 38% of the simulations, although the knot can move along the chain, it remains on the chain and is translocated through the pore after significant tightening (Fig. S5 in SI Appendix). The initial knot position is crucial to understand these differences. In such a case, the knot core is closer to the C-terminal for CTD-Tm1570 than NTD-TrmD (see Fig. 1C).

In the right panel in Fig. 5A, the degradation time distribution over 250 simulations at force equal 22 *ϵ/*Å is shown. According to the discussion above, NTD-TrmD and CTD-Tm1570 presenting different pattern in its degradation mechanism. The NTD-TrmD degradation time distribution is less widely spread, showing one pattern in its degradation mechanism. On the contrary, the degradation process for CTD-Tm1570 at force 22 *ϵ/*Å has shown different patterns. As a result, the degradation time distribution presents a multimodal behavior, which describes these multiple degradation mechanisms, translocation of the tight knot in different positions, and complete translocation when the knot slides off the chain.

Overall, our results stay in a reasonable agreement with the experimental data, the NTD-TrmD protein has been shown in both cases to be the protein with the fastest degradation. Meanwhile, CTD-Tm1570 has displayed a slow degradation. For the control, GFP protein has shown values between TrmD and Tm1570, in both cases, experimental and theoretical results.

A more comprehensive investigation of the degradation process of NTD-TrmD, GFP, and CTD-Tm1570 includes analyses related to the native contact map and the degree of local frustration. According to Fig. 1B, the knot core is closer to the C-terminal in CTD-Tm1570 than in NTD-TrmD. In Figures 5B and C, in the lower triangle, display the native contact map for NTD-TrmD and CTD-Tm1570, respectively. All performed simulations of NTD-

TrmD with force equal and less than 28 *ϵ/*Å showed that the knot slid completely off the chain in almost 100% of these simulations (Fig. S5 in SI Appendix). In contrast, analyses within the same range of forces for CTD-Tm1570 displayed different values that decreases from 98% at 28 *ϵ/*Åto only almost 20% at 22 *ϵ/*Å. The percentage numbers (%) correspond to the simulations with complete translocation of tight knot.

Due to the knot core position in the CTD-Tm1570 the number of native contacts that must be broken is much higher than in the case of NTD-TrmD. Detailed analysis reveals that it is more pronounced to observe different mechanisms during the degradation process for CTD-Tm1570 than in the case of NTD-TrmD. We must remember that all simulations were performed at low temperatures, therefore thermal fluctuations are not the main source of breaking local interactions, but the knot translocation plays an essential role during the degradation process at low forces.

Another source that could be responsible for and attributed to the difference in the degradation times for the target proteins is quantifying the degree of local frustration. Frustration can happen for geometric reasons or due to competition between interactions of the basic elements. In Figures 5B and C, upper triangle, we evaluate how many highly frustrated contacts are given in the target structures. Figures 5D and E display the distribution of highly frustrated contacts in the NTD-TrmD and CTD-Tm1570 native structures.

As discussed above, during the degradation process, the knot core can move and slide entirely off the chain towards the N-terminal. Based on the knot core position, the path is much longer for the knot translocation to CTD-Tm1570 than NTD-TrmD. Due to the number of native contacts that should be broken and the highly frustrated contacts, some kinetic traps could appear earlier than the complete translocation. There are several studies in which some kinetic traps are responsible for stabilizing intermediates states and changing the folding rate. The Im7 and Im9 proteins are one of the famous cases in which highly frustrated contacts are correlated to the stabilization of their intermediate states.^24^ Another case is related to the Interleukin (IL) family, in which the folding traps make IL-1*β* fold more slowly than IL-1Ra.^25^

Some notable examples in the literature describe the signals that encode proteins and correlate with minimally and highly frustrating interactions with their stability. In this study, using a minimalist model, we presented and highlighted from a macroscopic point of view the correlation between knot translocation, native contacts, and highly frustrating interaction. Even though the coarse-grained computational model may not be directly and quantitatively compared to the experimental results, it might be able to capture the essential features of the degradation process of the NTD-TrmD, GFP, and CTD-Tm1570 qualitatively.

## Discussion

In this study, we focus on the existence the first crystal double-knotted protein, *Cn*TrmD-Tm157, in which we focused on the degradation process from an experimental and theoretical perspective to understand how double and single-knotted protein might be degraded. Although knotted proteins may be vulnerable to degradation due to their complex structures, understanding the mechanisms by which cells degrade knotted proteins can provide insights into how cells maintain protein homeostasis and respond to stress or damage.

We investigated the degradation process of deeply double and single-knotted proteins, *Cn*TrmD-Tm1570, NTD-TrmD, and CTD-Tm1570, respectively, as well as unknotted protein, GFP, *via* ClpXP proteasome. The experimental results confirmed that double and single-knotted proteins can be degraded using the machinery present in a living cell (Fig. 3). Furthermore, the results for single knotted and unknotted proteins have shown a faster degradation for TrmD-like. Such a result was validated through direct comparison with another knotted protein from the same SPOUT superfamily, YbeA protein.

The main issue discussed and highlighted is related to the degradation process of the first double-knotted protein, *Cn*TrmD-Tm1570. According to the experimental results, SDS-PAGE gel, the *Cn*TrmD-Tm1570 may not be thoroughly degraded, and the same portion that remains from TrmD and Tm1570, additional bands, it could be observed for the individual single-knotted proteins (Fig. 3). The main reason for such an observation could be subject to several mechanical stress, such as stretching or twisting. Therefore, we constructed a coarse-grained model to understand the degradation process for double and single-knotted proteins.

The purpose of the theoretical representation was to examine the effectiveness of the coarse-grained degradation model, in which such results will be compared with experimental data. Another purpose was to examine whether differential effects depending on the force applied and the main reason for the observed kinetic degradation rate difference. As a result, the double-knotted protein has three different pathways by which the *Cn*TrmD-Tm1570 protein can be degraded. Complete degradation with knot translocation through the pore and degradation with one and both knots slipped off the chain. These observations called attention to different ways that *Cn*TrmD-Tm1570 may be degraded; on the other hand, a clear explanation for such mechanical stress remains unclear. However, additional advances are in progress on the theoretical model to better capture the essential features of the biological machinery.

In addition, the insight from degradation may conduct a clear explanation related to the folding process of double-knotted protein. Considering that degradation strongly depends on local conformational stability and unfolding cooperativity of the whole structure, here we observe and demonstrate how knots play a crucial role during the degradation process and how their position may modulate the kinetic rates changing degradation pathways. Once single-knotted proteins actively conduct biological function, this study will be generally applicable in further analysis to understand how they work in the fusion protein. Double-knotted proteins have been called attention due to their complex structures and the challenges involved in understanding them, stability and function. This study presented the first crystal structure of the double knotted protein, *Cn*TrmD-Tm1570; however, high-confidence proteins containing double knots are provided AlphaFold, and their structural analyses are deposited in AlphaKnot.^1^

Thus, understanding the structure and function of knotted proteins can provide important insights into how proteins work and how they can be regulated. Overall, it still needs to be fully understood how knotted protein affects degradation, and more research is needed to clarify the relationship between these two processes.

## Materials and Methods

### Protein Preparation

We conducted degradation experiments using ClpXP system for: Tm1570, TrmD, fusion of these two proteins, i.e. TrmD-Tm1570, and GFP. Tm1570 and TrmD proteins are homodimers with active centres located in knot region and have deep 3_1_-trefoil knot in their structure. Tm1570 protein has 193 amino acids and their knot core is located between 113 and 158 residues. TrmD protein from *Calditerrivibrio nitroreducens* has 243 residues and their knot core is located between 85 and 129 amino acids. Each of the the proteins was prepared as a substrate for ClpXP by adding an 11-amino acid ssrA tag to the C-terminus. Also, for purification by affinity chromatography, six HisTag amino acids were added to the N-terminus of each protein. To confirm that the knot is still present in proteins after the tags addition, size exclusion chromatography of each proteins was performed. Each proteins ran as a dimer (based on retention time), suggesting that it had assumed its native knotted state. For more details about protein preparation and purification see SI Appendix.

A truncated form of ClpX lacking the N-terminal domain (referred to as ClpXΔN_6_) was used in all experiments. Residues 1-62 of full-length *Escherichia coli* ClpX were deleted using PCR mutagenesis to construct ClpXΔN_6_. ClpXΔN_6_ and ClpP were expressed and purified.

### Densitometric Analysis and Michaelis Menten Determination

The intensity of the bands on the gel were determined by ImageJ software. The densitometric analysis was carried out with the use of specific corrections and was aimed at determining the rate of the degradation reaction. A detailed description can be found in the supplementary materials.

For TrmD and Tm1570 of the proteins for which a Michaelis-Menten curve was constructed, concentrations raging from 1.875 *µ*M to 20.0 *µ*M for Tm1570 and from 1.875 *µ*M to 53.5 *µ*M for TrmD and 1.875 *µ*M to 18.0 *µ*M for GFP were degraded in triplicate. The mean was then drawn from the obtained values of the reaction rate constant in Excel and the standard deviation was calculated. For the results obtained in this way, a Michaelis-Menten diagram was prepared, from which it was determined V_m*ax*_ and K_M_ parameters. The Michaelis-Menten plot was created by fitting the curve to the obtained results.

The progress of degradation process for TrmD-Tm1570 protein was observed at three different concentrations: 3.75 *µ*M, 7.5 *µ*M and 13.5 *µ*M. Prior to degradation experiment it was processed as a dimer in the size exclusion chromatography, suggesting that it had assumed its native state.

### Molecular Dynamics, topology detection

In order to provide a more detailed interpretation about the degradation process, we performed a molecular dynamics (MD) coarse-grained simulations based on the structure-based C*α* model (SBM-C*α*), as presented in details in ref. ^26^ The SBM-C*α* defines the potential in such a way that the minimum energy value is attributed to the protein’s native structure. Native contacts are mimicked by the Gaussian potential^27^ and it was determined by the shadow contact map algorithm^28^ with the standard parameters defined by the SMOG server.^29^ All the simulations were performed using Gromacs v4.5.4. ^30^ The topology, and knot core location of each structure selected from the simulation was determined using an implementation of the Alexander polynomial defined in the AlphaKnot database. ^1,31^

### Degradation Time and Tube Model

The protein degradation time is a parameter based on the Mean First Passage Time (MFPT) and the tube model was modeled based on the ref. ^32^ The full description of these approaches find in the Supporting Information - Theoretical Model and Molecular dynamics.

### Protein Frustration Analysis

The local frustration index were analyzed in the Protein Frustratometer 2 server. The local configurational frustration indices was calculated as described in ref.^33^

This work was supported by the National Science Centre [#UMO-2018/31/B/NZ1/04016 to JIS].

## Notes

### Competing Interest Statement

The authors have declared no competing interest.

